# The biphasic and age-dependent impact of Klotho on hallmarks of aging and skeletal muscle function

**DOI:** 10.1101/2020.07.22.207043

**Authors:** Abish Pius, Zachary Clemens, Sruthi Sivakumar, Amrita Sahu, Sunita Shinde, Hikaru Mamiya, Nathaniel Luketich, Jian Cui, Joerg D. Hoeck, Sebastian Kreuz, Michael Franti, Aaron Barchowsky, Fabrisia Ambrosio

## Abstract

Aging is accompanied by a disrupted information flow, which results from accumulation of molecular mistakes. These mistakes ultimately give rise to debilitating disorders such as skeletal muscle wasting, or sarcopenia. To estimate the growing “disorderliness” of the aging muscle system, we employed a statistical physics approach to estimate the state parameter, entropy, as a function of genes associated with hallmarks of aging. Although the most prominent structural and functional alterations were observed in the oldest old mice (27-29 months), we found that the escalating network entropy reached an inflection point at old age (22-24 months). To probe the potential for restoration of molecular “order” and reversal of the sarcopenic phenotype, we overexpressed the longevity protein, α-Klotho. Klotho overexpression modulated genes representing all hallmarks of aging in both old and oldest-old mice. However, whereas Klotho improved strength in old mice, intervention failed to induce a benefit beyond the entropic tipping point.

## INTRODUCTION

Aging is a universal process, and decades of research have gone into understanding the cellular mechanisms underlying aged tissue phenotypes. With the goal of conceptualizing common molecular and cellular mechanisms that underlie the effect of time’s arrow on mammalian tissue health, López-Otín et al. described nine “Hallmarks of Aging” (1). These hallmarks are: epigenetic alterations, cellular senescence, altered intercellular communication, telomere attrition, nutrient sensing deregulation, mitochondrial dysfunction, stem cell exhaustion, loss of proteostasis and genomic instability. A criterion in the identification of these hallmarks was that aggravation or attenuation of the biological process results in an accelerated aging or more youthful phenotype, respectively. These hallmarks, therefore, have the potential to pave the way towards the development of therapeutic approaches to counteract the effect of aging on organismal health.

Numerous longitudinal aging studies have positively associated skeletal muscle health with healthspan and longevity, and skeletal muscle strength has even been shown to be a reliable predictor of biological age and mortality (2-5). Furthermore, an age-related loss of skeletal muscle mass and function (i.e. sarcopenia) is associated with loss of mobility and increased fall risk (6-9). Tissue-level features of sarcopenia have been described extensively, and include a decreased myofiber size, increased intramuscular fat accumulation (myosteatosis), and a preferential atrophy of type II (fast-twitch) muscle fibers (10-12). However, our understanding of the cellular mechanisms underlying sarcopenia is still lacking, a shortcoming that has hindered the development of targeted and specific interventions (13-16). Currently, approaches to the treatment and prevention of sarcopenia largely focus on the prescription of physical activity and dietary modifications, strategies that have shown moderate success (17-19). Proven pharmacological interventions for sarcopenia do not exist, though some drugs such as vitamin D, insulin-like growth factors, and testosterone are in clinical trials (15, 20-23). Interestingly, many of the drugs currently being tested also act on pathways associated with the longevity protein, α-Klotho (Klotho).

Klotho is best known for its role in delaying age-related pathologies in various organ systems, including skeletal muscle (24-26). The pro-longevity effects of Klotho have been partially attributed to modulation of fibroblast growth factor 23 (FGF23), Wnt, and mTOR pathways (27, 28), several of which have also been a target in sarcopenia clinical trials (15, 22, 29). With age, circulating Klotho levels gradually decline, and epidemiological studies have revealed that decreased circulating Klotho levels are associated with an accelerated loss of skeletal muscle mass and strength (30-32). Similarly, mice deficient for Klotho display significant muscle wasting which is hypothesized to be caused by increased mitochondrial reactive oxygen species (ROS)(33, 34). These findings are consistent with the observation that mitochondrial accumulation of ROS leads to a decline in muscle mass and decreased regenerative potential (33, 35). Taken together, these studies suggest an unexplored mechanistic link between sarcopenia and a decline of Klotho with increasing age.

In this study, we first thoroughly characterize and compare structural, functional, and genomic changes in skeletal muscle across the lifespan in mice. In order to derive a global metric of the loss of molecular fidelity over time, we used an information-based calculation of network entropy. Changes in the overall gene expression profile were captured using protein-protein interaction (PPI) networks. A higher network entropy means a probabilistically greater degree of disorganization or randomness in the system. We found that network entropy increases from young (4-6 months) to old (21-24 months) mice, but then plateaus in the oldest-old mice (27-29 months). We next tested the ability of AAV-mediated Klotho delivery (AAV-Kl) to attenuate these changes and counteract sarcopenia in old and oldest-old mice. Whereas old mice displayed a signficant improvement in function, AAV-Kl failed to induce a benefit in the oldest-old mice. Furthermore, AAV-Kl regulated a multitude of genes associated with all hallmarks of aging. Unexpectedly, however, the direction of change in response to treatment was, for many genes, age-dependent. Taken together, these results suggest Klotho enhances skeletal muscle structure and function at the early stages of sarcopenia, but the beneficial effects are lost after the entropic inflection point.

## RESULTS

### Sarcopenic changes are subtle until mice reach an advanced age

To thoroughly describe the trajectory of sarcopenic alterations according to sex and over time, we characterized muscle structure and function in young, middle-aged, old, and oldest-old male and female C57Bl/6J mice. Mouse age groups were selected to parallel stratifications commonly used in epidemiological studies and correspond, roughly, to individuals aged 20-30 (36), 38-47, 56-69, and 78+ years (37, 38). As expected, a progressive decline in tibialis anterior (TA) wet weight (normalized to body weight) was observed over time for both males and females, though the percent decrease at the oldest-old age was significantly greater in male mice (34%) when compared to females (17%; **Figure 1A**). A decline in muscle weight was concomitant with a decline in physiological cross-sectional area (CSA), calculated as 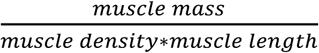 (39)(**Figure 1B**), though there were only minor differences in mean myofiber cross-sectional area across age groups as determined by histological analysis (**Figure 1C, D**).

**Figure 1.**
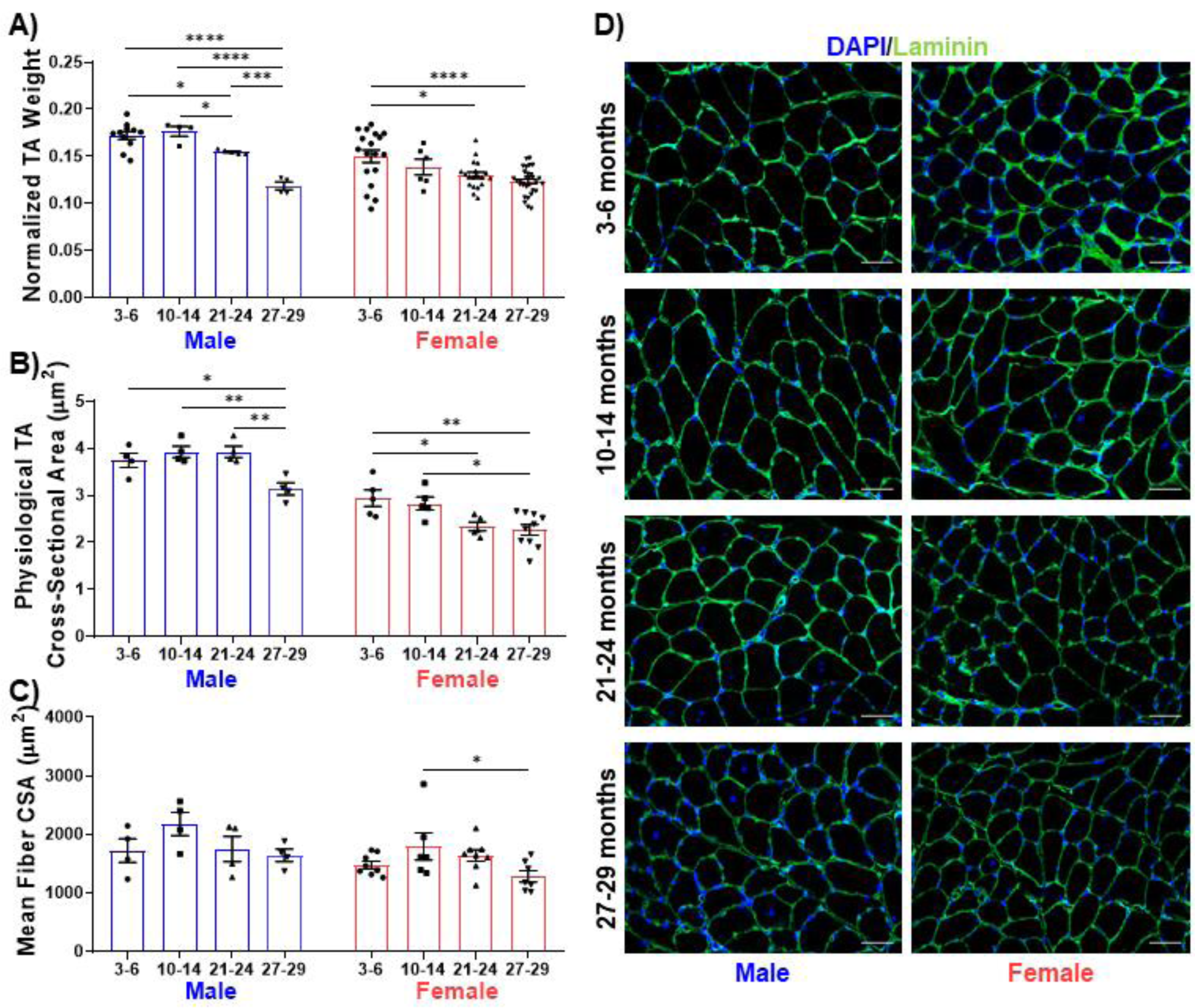
Declines in muscle structure are subtle until advanced age. (A) Tibialis anterior (TA) muscle weight as a percentage of whole-body weight in young (3-6 months), middle-aged (10-14), old (21-24), and oldest-old (27-29) male (n = 4-11/group) and female mice (n = 6-18/group) (*p<.05, ***p<.001, ****p<.0001, one-way ANOVA). (B) Mean physiological cross-sectional area of the TA muscle of male (n = 4/group) and female mice (n = 5-10/group) (*p<.05, **p<.01, One-Way ANOVA). (C) Average fiber cross-sectional area of uninjured male (n = 4/group) and female (n = 6-8/group) mice across age groups (*p<.05, one-way ANOVA). (D) Representative images of histological sections stained for laminin. Scale bars = 50 µm. All data presented as mean ± SEM error bars.

Whole-body muscle strength, as measured by the four-limb hang test, gradually declined with increasing age for both male and female mice (**Figure 2A**). However, *in situ* contractile testing of the TA muscle revealed a significant decline in muscle strength only in the oldest-old mice, regardless of sex (**Figure 2B, C, D**). Changes in temporal characteristics of the muscle contraction profile, including increased time to peak twitch contraction and half relaxation time, were most prominent in the oldest-old female group (**Figure 2E, F)**. These changes are consistent with previous reports of preferential atrophy of fast-twitch (type II) fibers with increased age (12, 40, 41). Taken together, these data suggest that old mice display a mild sarcopenic profile and that common features associated with clinical metrics of sarcopenia only become evident in oldest-old mice.

**Figure 2.**
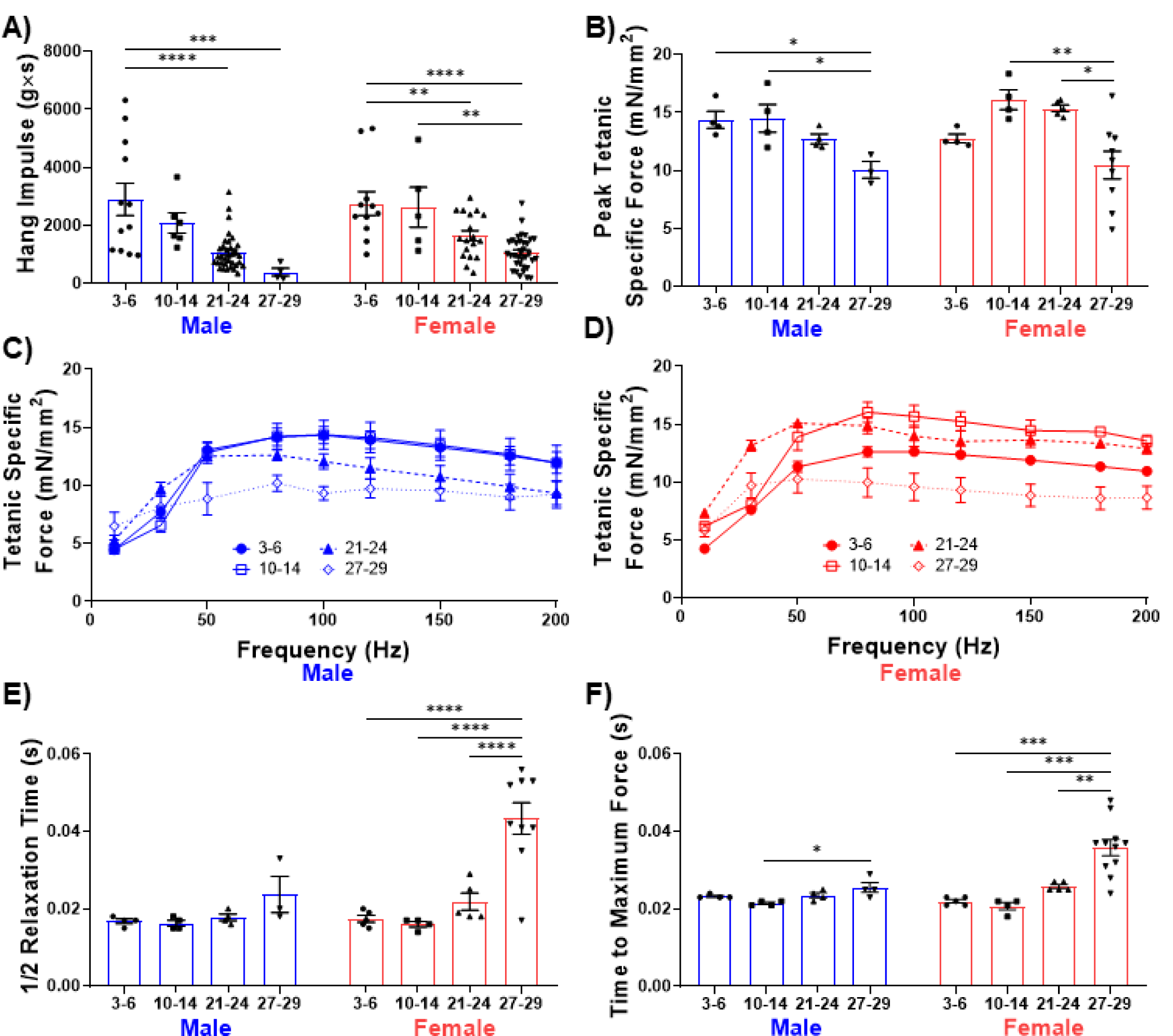
Male and female mice display a progressive loss of muscle function over time. (A) Whole-body endurance of male (n =4-35/group) and female (n = 5-34/group) mice measured via the four-limb hang test (**p<.01, ***p<.001, ****p<.0001, One-Way ANOVA). (B) Peak specific tetanic force production in male (n=3-4/group) and female (n = 4/9group) TA muscles (*p<.05, **p<.01, One-Way ANOVA). (C) Force frequency curves for male mice (n = 3-4/group). (D) Force frequency curves for male mice (n = 4-9/group) mice. (E) Half relaxation time following single twitch stimulation (****p<.0001, One-Way ANOVA, male mice (3-4/group), female mice (4-9/group)). (F) Time to maximum force following single twitch stimulation (*p<.05, **p<.01, ***p<.001, One-Way ANOVA, male mice (3-4/group), female mice (4-9/group)). All data presented as mean ± SEM error bars.

### Aging drives a progressive disruption in genes associated with hallmarks of aging

Our structural and functional findings across age groups revealed exaggerated changes between old and oldest-old female mice. We next used RNA-Seq analysis to investigate changes in the transcriptome of young, old, and oldest-old female hindlimb muscle (**Figure 3A**). We excluded middle-aged muscles from the analyses since this age group was phenotypically similar to old mice. Principal Component Analysis (PCA) revealed distinct gene expression profiles according to age, with young muscle segregating from the old and oldest-old muscle profiles, the latter of which slightly overlapped (**Figure 3B**). This is consistent with previous transcriptomic studies of human skeletal muscle in which the bulk of gene expression changes plateaued over time (42, 43).

**Figure 3.**
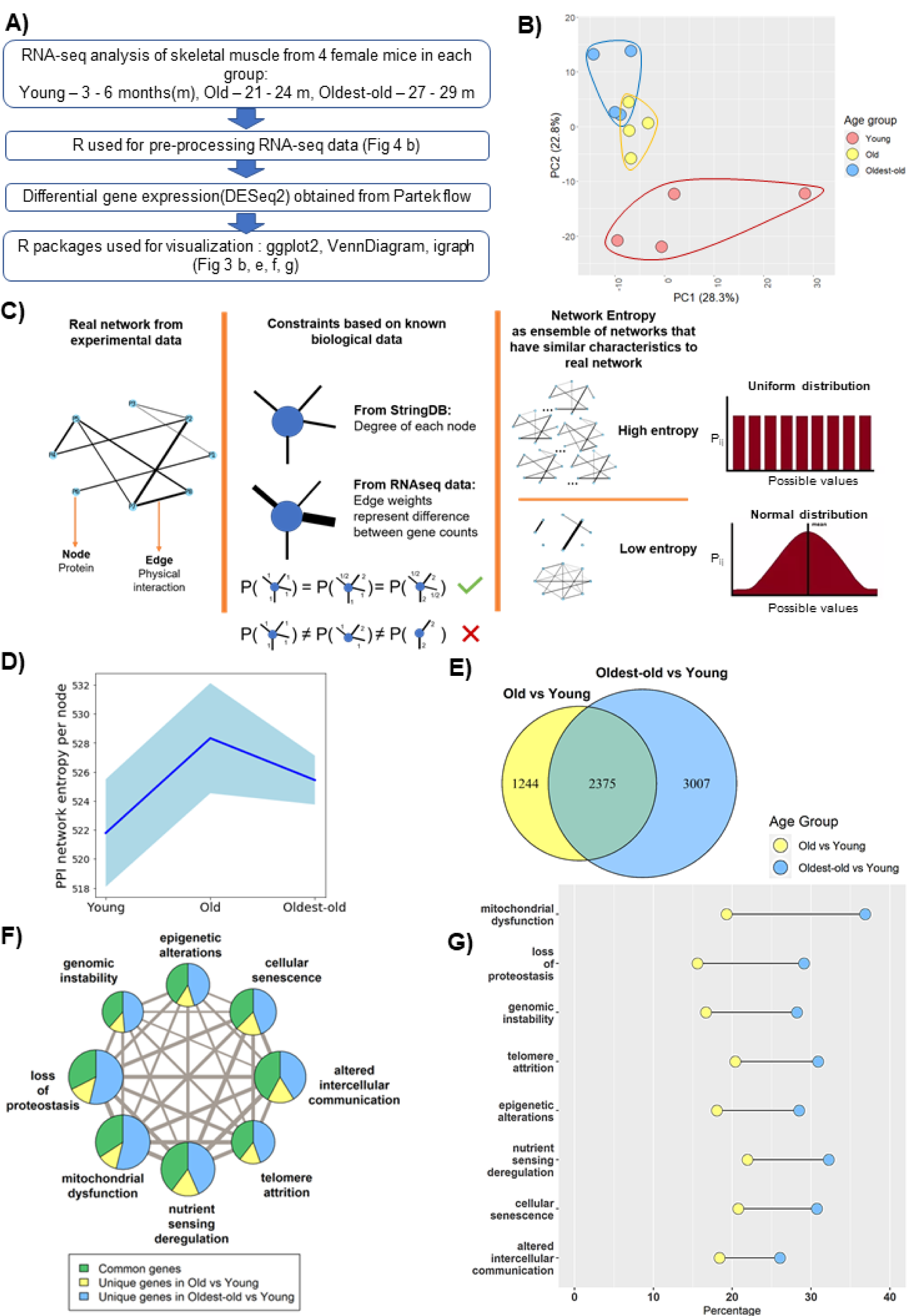
Network entropy increases from young to old mice, after which time it plateaus. (A) RNA-seq analysis workflow. (B) Principle component analysis (PCA) showing overall gene expression patterns in young, old, and oldest-old mice (n = 12). (C) Schematic description of network entropy computation and interpretation. PPI networks were generated based on RNAseq data. We capture the degree sequence and the edge weights from the network obtained from experimental data in the form of constraints. The ensemble of networks that follow these constraints have similar network features. If the probability distribution is skewed, then it has a low network entropy and if not, then it is has high network entropy. (D) Protein-protein interaction (PPI) network entropy computed from transcriptomic data indicates an increase in transcriptional noise of hallmarks of aging genes. A non-parametric Kruskal Wallis test (p=0.0741) and Dunn’s post-hoc test was performed. Entropy of young to old changed with p=0.07. (E) Venn Diagram showing the number of differentially expressed (DE) genes between groups Old vs Young, and Oldest-old vs Young mice. (F) Network plot denotes the total number of DE genes per hallmark, and the corresponding proportion of of DE genes that are unique to old and oldest old, when compared to young counterparts for each hallmark of aging. Edge weights denote the number of genes that are common between the two hallmarks the edge connects. The node sizes are proportional to the number of genes that fall into each hallmark. (G) Bar plot showing the ratio of DE genes between old and oldest old groups within each hallmark of aging, when compared to young counterparts.

To date, most RNA-seq studies in skeletal muscle have used a bottom-up approach, focusing on differentially expressed (DE) genes and their corresponding pathways. While this approach is valuable for identifying specific signaling pathways that may contribute to sarcopenia, it does not give a holistic view of the disease mechanism. We therefore employed a top-down approach and focused on shared features of age-related decline (1). This is consistent with Hayflick’s proposition that “the common denominator that underlies all modern theories of aging is change in molecular structure and, hence, function,” and, as such, the ultimate cause of an aged phenotype is “increasing molecular disorder”, or entropy (44). Although the entropy theory of aging has existed for centuries, it is has generally been considered in the context of thermodynamics (i.e. “dissipation” of energy at the biomolecular level). While theoretically intriguing, this form of entropy is challenging to quantify. Given that the essence of entropy is a loss of information in a system, we estimated network entropy as described by Menichetti et al (45). To do this, protein-protein interaction (PPI) networks were generated from RNAseq data across young, old and oldest-old age groups. Given our focus on age-related alterations, only those genes associated with the hallmarks of aging were included in the analysis. In the network, nodes represent the corresponding proteins, edges represent the physical interaction of proteins, and edge weights represent the differential gene expression (**Figure 3C**). Genes associated with “stem-cell exhaustion” largely overlapped with all other hallmarks, and, hence, were excluded from our analysis **(Supplemental methods)**.

Consistent with PCA findings, but contrary to phenotypic changes, we observed the greatest increase in network entropy between young and old groups, with little difference between the old and the oldest-old (**Figure 3D**). This suggests that gene expression changes likely precede the observed functional deficits in oldest-old mice (**Figure 2E, F**). The same entropy trend emerged even when all genes were considered, suggesting that the subset of genes associated with the hallmarks of aging capture the overall tissue transcriptomic profile without loss of information (**Supplemental Figure 1**). Furthermore, we found that the increase in entropy between young and old muscles was driven by a relatively small pool of genes that displayed a large magnitude of change (**Supplemental Figure 2**). In contrast, there was a much larger number of DE genes between old and oldest-old muscle, though the magnitudes of change were generally low (**Supplemental Figure 2**). This suggests a loss of specific changes and a broadening of gene expression alterations across the genome in the oldest-old age group.

We next probed for “weakest links” among the hallmarks of aging in which the effect of aging may become manifest prior to the others. Unexpectedly, we found that all of the hallmarks were similarly affected when considering changes between young and old mice and young and the oldest-old mice, though roughly twice as many genes changed in the oldest-old mice (**Figure 3 E,F,G**). This further implies a dysregulation of the transcriptomic profile in the oldest-old mice, a finding that is consistent with previous reports (42).

### AAV-Klotho administration improves muscle regeneration in old mice

Given that a criterion in the identification of aging hallmarks is a susceptibility to interventions that may aggravate or attenuate age-related declines, we next tested whether the declines in muscle function and structure observed in aged mice could be attenuated through Klotho supplementation. As a first step, we confirmed that circulating Klotho indeed declines with age in mice, consistent with human findings (30) (**Figure 4A**). In both sexes, we also observed a stepwise increase in circulating fibroblast growth factor-23 (FGF23), a known co-factor for Klotho signaling and a key hormone in the regulation of mineral ions and vitamin D (46) (**Figure 4B**).

**Figure 4.**
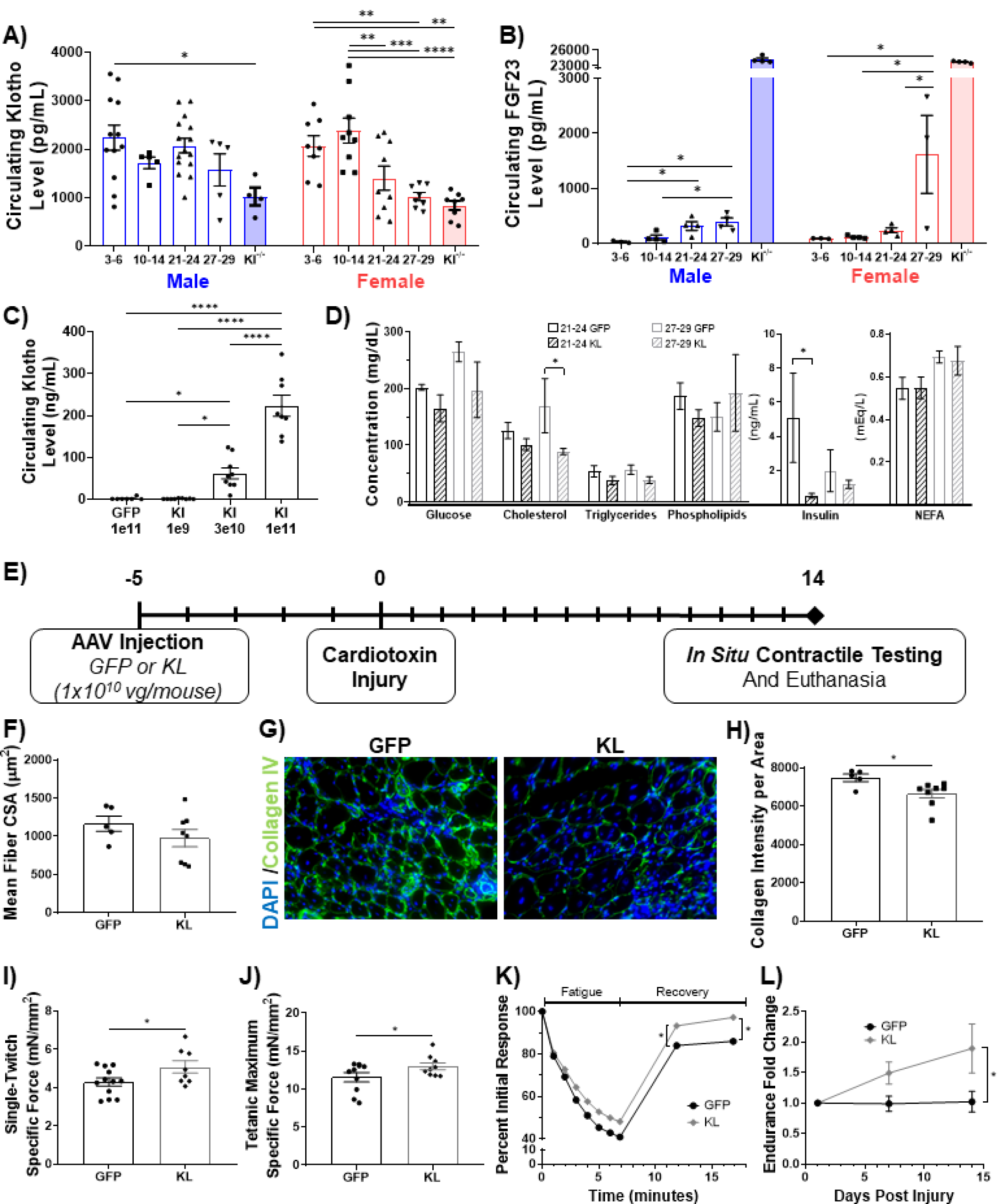
Gene delivery of Klotho enhances functional muscle regeneration following an acute injury. (A) Changes in circulating Klotho levels measured via ELISA (*p<.05, **p<.01,***p<.001, ****p<.0001, one-way ANOVA, male n = 4-15/group, female n = 8-9/group, *Kl-/-*negative control). (B) Changes in circulating FGF23 levels (*p<.05, **p<.01, One-Way ANOVA, KO values were excluded from statistical analysis, male n = 3-4/group, female n = 3/4/group, *Kl-/-*control). (C) Circulating Klotho levels in young male (n = 7-9/group) mice injected with AAV-Klotho at varying doses. (D) Serum metabolite concentration levels for AAV-GFP(GFP)- and AAV-Kl (KL)-treated female mice (*p<.05, One-tailed Student’s t-test, n = 14, female mice(3-5/group)). (E) Experimental design schematic. (F) Histological quantification of myofiber cross-sectional area (n = 5-8/group). (G) Representative images of injured tibialis anterior muscles stained for collagen IV (green) and DAPI (blue). (H) Collagen IV expression in GFP-versus KL-treated mice (*p<.05, one-tailed student’s t-test, n = 5-8/group). (I) TA specific twitch force produced 14 days post injury (dpi) (*p<.05, one-tailed Student’s t-test, n = 8-12/group). (J) TA peak tetanic specific force 14 dpi (*p<.01, one-tailed Student’s t-test, n = 9-10/group). (K) Change in force production of the TA over time as mice underwent a fatigue protocol consisting of repeated TA stimulation for a total of 7 minutes followed by recovery over two five-minute intervals (*p<.05, two-way ANOVA, n = 8-10/group). (L) Fold change in whole body endurance compared to 1 dpi hang impulse score (*p<.05, One-tailed Student’s t-test, n=7-9/group). All data presented as mean ± SEM error bars.

An AAV vector carrying a mouse Klotho full-length cDNA (AAV-Kl) was constructed and administered to old mice via tail vein injection **(Supplemental figure 3)**. Meso Scale Discovery-Enzyme-linked Immune Sorbent Assay (MSD-ELISA) conducted on serum collected 3-weeks after the administration confirmed a dose-dependent increase in circulating Klotho levels (**Figure 4C**). The AAV expression of Klotho had no significant effects on non-fasting serum metabolites, although AAV-Kl muscles had a slight, but not significant, decrease in cholesterol and insulin (**Figure 4D**). Next, to confirm the physiological effect of AAV-Kl administration, we tested whether AAV-Kl administration could replicate data from a previous report demonstrating improved regeneration following supplementation with recombinant Klotho protein (33). AAV-Kl (1×10^10^ vector genomes/animal) or an equal titer AAV-GFP control was administered via tail vein injections to old male mice five days before injuries to the tibialis anterior (TA) muscles bilaterally with cardiotoxin (**Figure 4E)**. There was no difference in myofiber cross-sectional area in the muscles of animals treated with AAV-Kl when compared to control counterparts (**Figure 4F**). However, fibrosis was significantly decreased in AAV-Kl-treated mice, as evidenced by decreased collagen IV abundance (**Figure 4G, H**). Mice receiving AAV-Kl also showed increased specific twitch and peak specific tetanic force production (**Figure 4I, J**). Resistance to a fatiguing protocol was not significantly affected by AAV-Kl treatment (**Figure 4K**). However, force recovery following completion of the fatiguing protocol was significantly enhanced with AAV-Kl (**Figure 4K**). Additionally, whole-body strength improved almost two-fold over 14 days for mice treated with AAV-Kl, while control mice showed no improvement (**Figure 4L**). These results confirmed that elevated Klotho levels systemically enhance skeletal muscle regeneration, similar to previous reports using intraperitoneal injection of recombinant Klotho (33, 47).

### Intravenous AAV-Klotho administration enhances muscle structure and function in old, but not in the oldest-old, mice

Next, we tested whether AAV-Kl administration enhances muscle function in uninjured mice. Old and oldest-old mice received either AAV-Kl or AAV-GFP at a titer of 3×10^8^vg via tail vein injection. This dose was chosen because it provided the greatest improvement in muscle strength, as determined over the course of a preliminary dose-testing study performed in old mice (**Supplemental Figure 3**). A challenge of studying sarcopenia in mouse models is the high rate of incidental mortality and the onset of confounding pathologies in very old animals. Of the 44 old and oldest-old mice included, only 23 mice were included in final analyses due to incidental morbidity and mortality (**Figure 5A**).

**Figure 5.**
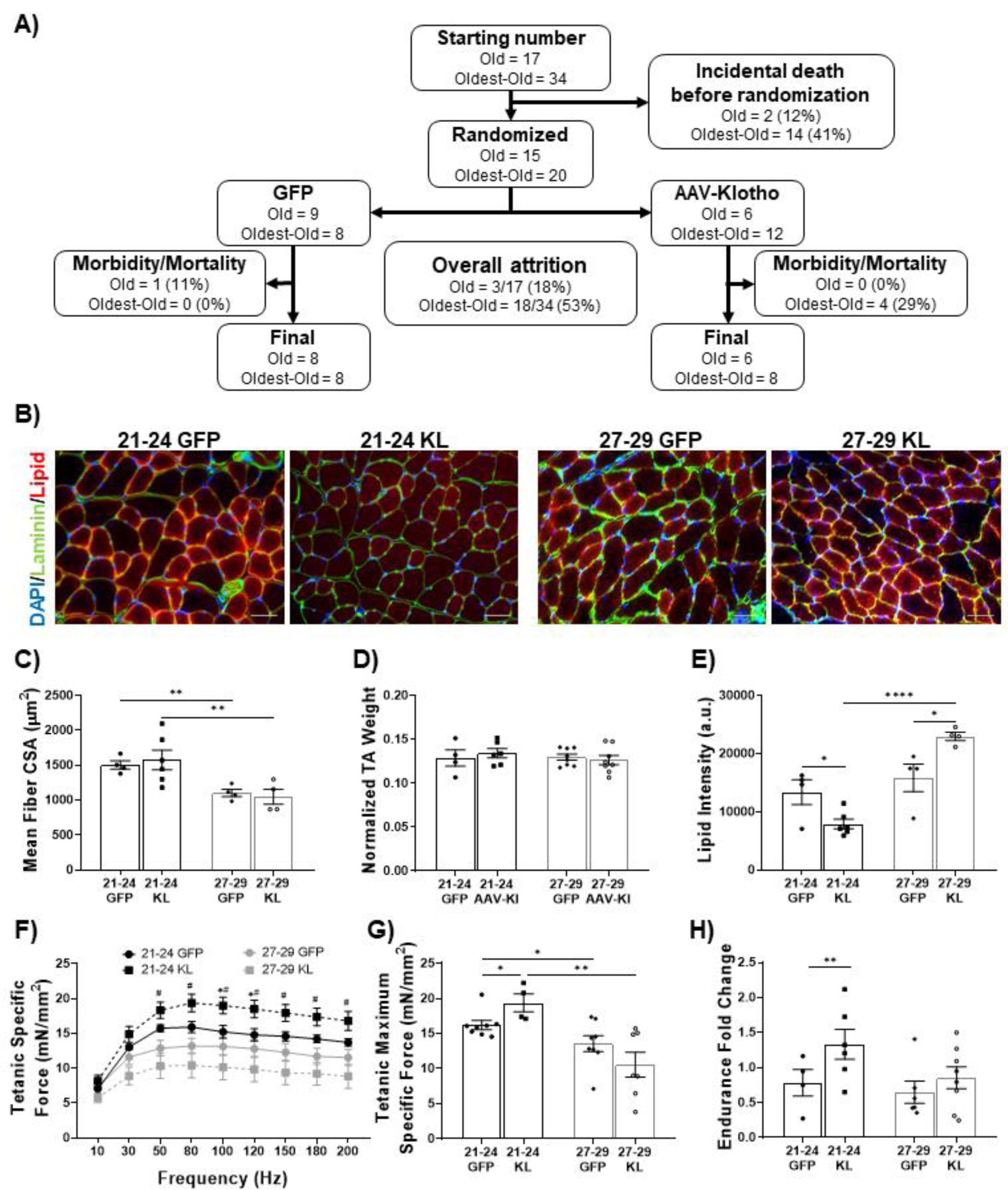
AAV-Klotho enhances muscle function in old, but not oldest-old, mice. (A) Animal inclusion flow chart. Mortality describes mice that died over the course of the experiment. Morbidity describes mice whom pathology was found in at euthanasia via necropsy and were therefore excluded from analyses. (B) Representative images showing myofiber area (Laminin; green), lipid (red), and DAPI (blue) of the TA in old and oldest-old female mice treated with GFP or Kl. Scale bars = 50 µm. (C) Quantification of muscle-fiber cross-sectional area (**p<.01, one-tailed Student’s t-tests, n = 4-6/group). (D) TA wet weight normalized to body-weight (n=4-8/group). (E) Lipid intensity in cross-sections of the TA (*p<.05, ****p<.0001, one-tailed Student’s t-tests, n = 4-6/group). (F) Force-frequency curve for all experimental groups with tetanic stimulation (*p<0.05 comparing 21-24 KL group to age matched controls, #p<0.05 comparing 27-29 KL group to age matched, two-way ANOVA repeated measures (n=4-6/group). (G) TA peak specific force production under tetanic stimulation (*p<.05, **p<.01, one-tailed Student’s t-tests, n = 4-8/group). (H) Hang-test performance 14-days after injection of AAV-Kl or AAV-GFP, calculated relative to baseline performance (**p<.01, one-tailed Student’s t-test, n = 4-8/group). All data presented as mean ± SEM error bars.

Histological analysis of AAV-Kl-treated old and oldest-old mice revealed no difference in the myofiber cross-sectional area or muscle weight when compared to controls (**Figure 5B-D)**, but did significantly reduce lipid accumulation in the skeletal muscle of old mice when compared to age-matched controls (**Figure 5B, E**). In contrast, oldest-old mice receiving AAV-Kl displayed higher lipid abundance when compared to age-matched controls (**Figure 5E**). These findings were consistent with functional metrics, in which maximum force production improved with AAV-KL treatment in the old, but not the oldest-old group (**Figure 5F, G**). Accordingly, whole-body endurance of old mice receiving AAV-Kl was 30% greater 14 days after injection when compared to pre-injection endurance, while oldest-old mice showed no improvement with AAV-Kl administration (**Figure 5H**). These findings suggest that systemic upregulation of Klotho induces therapeutic benefits in old mice, but not in mice of more advanced age with a more pronounced sarcopenic profile.

### Klotho affects hallmark of aging genes across age groups, but oldest-old mice exhibit a dysregulated response

To understand the benefit of Klotho in the old group and lack of benefit in the oldest-old group, we revisited the transcriptomic profiles across groups according to the hallmarks of aging. When comparing age matched control and AAV-Kl treatment groups, we observed approximately three times more DE genes in the oldest-old group treated with Klotho as compared to the old group (**Figure 6A**). Existing literature suggests that Klotho attenuates mitochondrial dysfunction, telomere attrition, and cellular senescence (33, 48-50). However, we found that AAV-Kl affected all the hallmarks similarly (**Figure 6B, C**).

**Figure 6.**
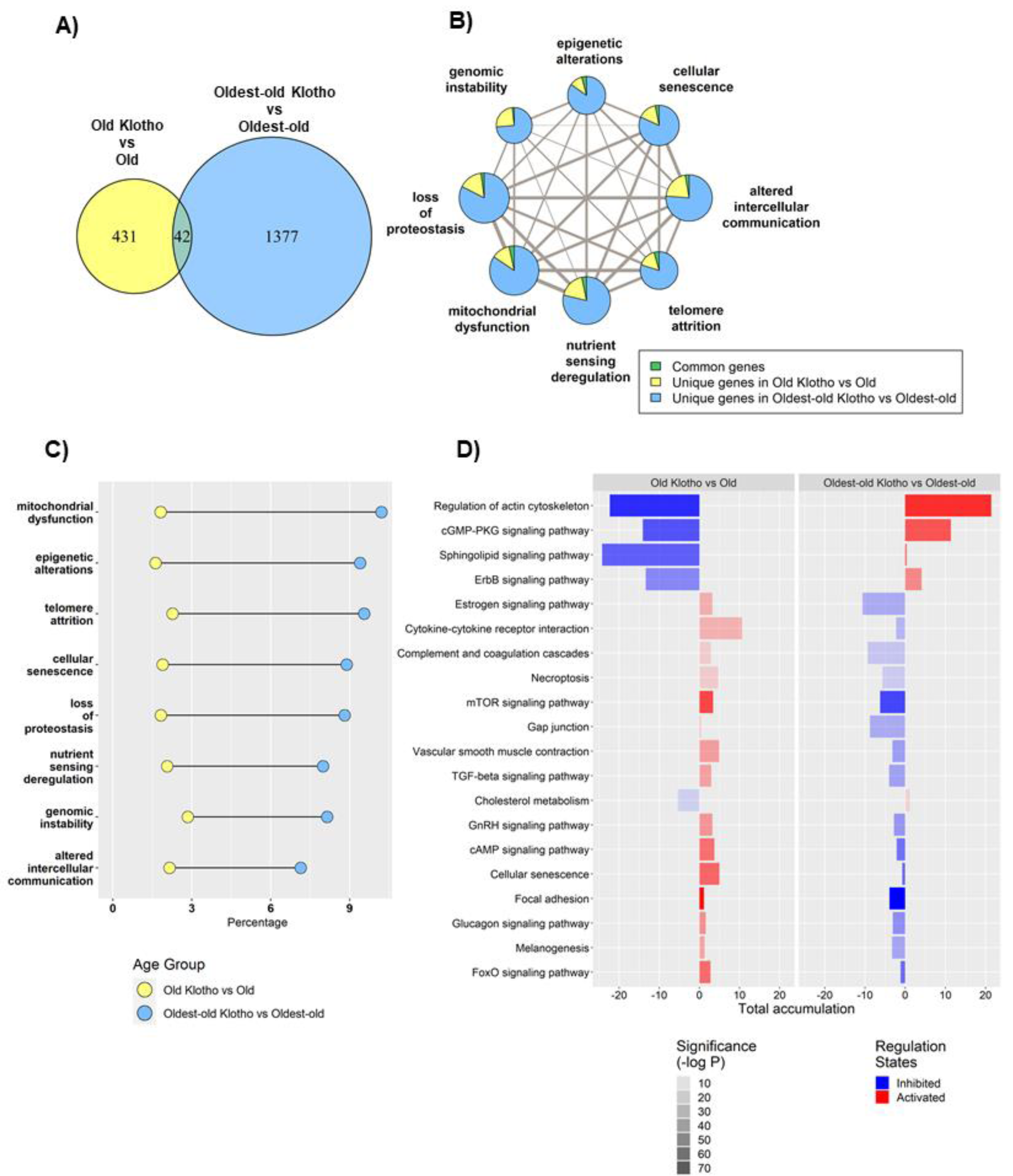
The effect of AAV-Kl administration on genes associated with hallmarks of aging is age-dependent. (A) Venn Diagram showing the number of differentially expressed (DE) genes between groups treated with AAV-Klotho (n = 4) vs AAV-GFP (n = 4) mice. (B) Network plot with each node as a pie chart that denotes the total number of DE genes in that hallmark, and the wedges denote the proportion of DE genes between groups treated with AAV-Kl vs AAV-GFP for each hallmark of aging. The edge weights denote the number of genes that are common between the two connected hallmarks. The node sizes are proportional to the number of genes that fall into each hallmark. (C) Bar plot showing the ratio of DE genes between groups treated with AAV-Kl vs AAV-GFP within each hallmark of aging. (D) This bar plot shows top 20 KEGG pathways that change differently between old and oldest-old groups after AAV-Kl treatment ranked by largest absolute difference in total accumulation.

Next, we performed Kyoto Encyclopedia of Genes and Genomes (KEGG) pathway enrichment analysis to identify pathways that had contrasting responses to Klotho treatment between the old and the oldest-old groups. Regulation of actin cytoskeleton, cGMP-PKG signaling pathway, and sphingolipid signaling pathway were among the top pathways in which the gene response to Klotho was opposed in the old versus oldest-old mice (**Figure 6D**). These findings suggest that Klotho’s response in oldest-old mice may disrupt pathways related to cell structure, cell membrane integrity, and intercellular communication, which is consistent with the increased myosteatosis and decreased contractile capacity of oldest-old mice treated with Klotho when compared to old counterparts (**Figure 5B, E, F, G, Supplemental figure 5)**.

## DISCUSSION

In this study, we evaluated phenotypic and gene expression trajectories across the lifespan of mice. Whereas declines in structure and function were observed in old mice and more prominently in oldest-old mice, alterations in the expression of genes linked to the hallmarks of aging displayed a non-linear trajectory, with the most significant changes in gene expression profiles occurring between young and old mice. Furthermore, we found that AAV-Kl supplementation improved skeletal muscle vitality in the old, but not the oldest-old, mice. Notably, genes linked to mitochondrial functioning, lipid signaling, and cell structure appeared most dysregulated in oldest-old mice following Klotho treatment, suggesting that regulation of metabolic pathways may be compromised in oldest-old mice following Klotho intervention.

In humans, sarcopenic declines in skeletal muscle mass typically manifest in the fourth decade of life (51, 52). The onset of sarcopenia in mice, on the other hand, has not been well-documented (53, 54), and studies of sarcopenia in animal models often exclude the oldest-old cohort (53, 55, 56). We found that decreased muscle mass and function – two of the classical markers of sarcopenia – become prominent in the oldest-old group for both male and female mice, whereas muscle declines in old mice are relatively modest. Additionally, the onset of sarcopenia in mice appears to be sexually dimorphic, with females displaying slightly accelerated functional declines. These findings are in agreement with one previous clinical report suggesting that sarcopenia progresses more rapidly in females (57), though other studies demonstrated minimal differences between males and females (11, 58). Future studies are needed to elucidate potential mechanisms by which the progression and severity of sarcopenia may be mediated by sex.

Clinical studies performed in older adults have suggested that loss of muscle function is predominantly a result of a loss of muscle quality, as opposed to a loss of muscle mass (9). Likewise, in our study, age-related functional declines were not accompanied by a significant loss of muscle mass. Instead, muscle mass and myofiber area were, for the most part, preserved over time. Although muscle mass decreased by only approximately 10% between the old and oldest-old age groups, the accompanying decrease in peak force production was over two-fold greater. The data suggests that poor muscle quality, such as the accumulation of intramuscular lipid, may be a more potent contributor to functional declines over time. Myosteatosis reduces muscle strength and blunts muscular activation in response to exercise (59), and it has been proposed that myosteatosis drives declines in function through impairment of muscle oxidative capacity, causing fiber-type transition towards slow-twitch fibers (60, 61). Indeed, temporal features of muscle contractile characteristics reveal that half relaxation time and time to maximum force were significantly increased in our oldest-old mice, consistent with a sarcopenic shift towards slow-twitch fibers (41). The fiber-type shift may also be attributed to decreasing activity levels with increasing age (9, 41).

In order to describe how changes in gene expression may relate to the phenotypic changes observed with aging, we implemented an information-based approach to calculate network entropy. Though an aging system is not necessarily closed (a now overturned criteria for an entropic system (44),(62)), focusing on the interconnectedness of network level protein-protein interactions allows for a single, integrative metric to estimate the “disorderliness” of the system (45). In our study, “disorderliness” of the gene expression profiles over time is represented by the probability of a network connection, mathematically similar to Shannon entropy (63). Similar to a previous report using microarray gene data from human circulating T-lymphocytes, we observed that network entropy increased from young to old mice (45). The effect size for network entropy in our sample is also comparable with this previous report (45). However, whereas T-lymphocytes displayed a sharp decline in the oldest-old group, potentially suggestive of a “survival effect” (45), we observed a plateau in entropy after old age (44, 45, 62). The data suggest that this peak then plateau may result from a small subset of genes that display highly specific changes from young to old age, followed by a large number of small, non-specific changes between old and oldest-old ages. Our observation that network entropy peaks at old age but that phenotypic changes are not prominent until later in life, supports the hypothesis that gene expression changes precede phenotypic changes (50). Evaluation of entropy at additional time points across the lifespan would be valuable to better identify an optimal therapeutic window. A therapeutic challenge, then, lies in how to promote healthier aging and best address the root of these gene expression changes before a sarcopenic phenotype develops. Moreover, though here we focused on quantification of entropy at the gene-level, given the progressive loss of histological and functional muscle integrity, it would be interesting to determine whether entropy at the proteomic level (e.g. using mass spectrometry) is able to capture entropic changes even into very old age.

Given the aforementioned observations, we wondered whether modulation of genes associated with the hallmarks of aging is capable of restoring a more youthful skeletal muscle profile. Indeed, AAV-Kl administration modulated all hallmarks of aging to comparable extents, though the magnitude of change was manifold times greater in the oldest-old mice. This enhanced response in oldest-old mice was not, however, accompanied by a greater therapeutic benefit. Instead, we observed a significant increase in muscle function in old mice receiving AAV-Kl, but intervention failed to produce a benefit in the oldest-old mice. Similar observations of treatment resistance have been reported in advanced age and critically ill populations in other pharmaceutical treatment studies (64, 65). In humans, dieting and exercise intervention studies reveal blunted muscle protein synthesis in the elderly population (greater than 60 years old) (66, 67). The lack of beneficial response to treatment is hypothesized to stem from deficiencies in downstream anabolic pathways, i.e. ‘anabolic resistance’ (65, 68, 69). In order to identify potentially compromised signaling pathways in oldest-old mice that may preclude a therapeutic benefit of AAV-Kl administration, we compared KEGG pathway activation differences across treatment groups. Old and oldest-old mice displayed opposing responses to AAV-Kl in pathways associated with cellular structure, lipid membrane integrity, and intercellular communication. DE in the old animals treated with AAV-Kl were primarily associated with paracrine signaling molecules such as F2 and Kng2, when compared to age-matched control counterparts **(Supplemental Table 2**). Intriguingly, both of these genes are associated with pathways playing a role in angiogenesis and vascular permeability (70, 71). These findings raise the possibility that restoration of vascular supply may be important in the development of therapeutics for sarcopenia. Indeed, Klotho has been shown to exert pro-angiogenic effects in several tissues, including the heart and skin (72, 73). On the other hand, AAV-Kl in the oldest-old mice modulated expression of numerous genes, the bulk of which were unrelated to vascularity. F2 and Kng2 were not responsive to AAV-Kl in this age group. These findings suggest that Klotho overexpression may have stimulated a haphazard transcriptional response in the oldest-old mice **(Supplemental Table 2)**. Furthermore, the sphingolipid signalling pathway was inhibited in old but not in the oldest-old, following Klotho treatment. Previous aging studies in *C*.*elegans* and *D.melanogaster*, suggest that inhibition of sphingolipid synthesis may be beneficial for maintaining proper lipid homeostasis and even extending lifespan (74, 75). This could explain our observation of increased lipid accumulation in the oldest-old following Klotho treatment, though further investigation is warranted.

Taken together, the data presented here suggest that intervention with AAV-Kl may be more effective in slowing the progression of sarcopenia at an earlier timepoint, rather than rescuing advanced pathology, at which time the transcriptomic response to intervention appears to be more stochastic. An interesting area of future investigation includes the determination of whether network entropy and PPI network architecture may be predictive of the efficacy of therapies designed to counteract the effect of time on skeletal muscle health and function.

## METHODS

### Animals

All animal experiments were approved by the University of Pittsburgh’s Institutional Animal Care and Use Committee. Experiments were performed using young (3-6 months), middle-aged (10-14 months), old (21-24 months), and oldest-old (27-29 months) male and female C57 BL/6J mice obtained from Jackson Laboratories. Prior to inclusion in experiments, animals were evaluated and those with visible health abnormalities were excluded. Given the large amount of variation in physical endurance capacity, animals were randomized into cohorts only if they met the criteria of falling into the 25^th^-75^th^ percentile for the four-limb hang test (described below). Each experiment was repeated across a minimum of two cohorts, and experimenters performing endpoint analysis were blinded to the experimental group.

### Immunofluorescence Staining, Imaging, and Analysis

Freshly harvested TA muscles were frozen by submersing them in frigid 2-methylbutane (cooled using liquid nitrogen) for one minute. Frozen tissues were sliced into 10µ sections. Antibody staining for tissue sections was conducted following these basic steps: 2% PFA in PBS solution for 10 minutes, PBS wash, .1% Triton-X in PBS for 15 minutes, .1% Triton-X+3% BSA in PBS for 1 hour, primary antibodies overnight at 4°C, wash, secondary antibodies in antibody solution for 1 hour, wash, DAPI (1:500 in PBS) for 2 minutes, wash. Primary antibodies used are shown in the table below.

| Antibody | Source | Dilution (in .1% Triton-X + 3% BSA+ 5% goat serum in PBS) |
| --- | --- | --- |
| Rabbit anti-Laminin | Abcam ab11575 | 1:500 |
| Lipidtox | Invitrogen H34476 | 1:200 |
| Collagen IV | Abcam ab6586 | 1:500 |

Secondary antibodies were selected based on the species of their primary antibody target and diluted 1:500 in the same solution as primaries. For sections from our AAV-treated animals, no secondary antibodies fluorescing in the green spectrum were used in order to avoid interference from GFP expression of the NTCs. Coverslips were mounted using gelvatol, then dried and imaged using a Nikon Observer Z1 microscope at 20x or 40x. Myofiber area was calculated by manually measuring the laminin rings using ImageJ software. Lipidtox intensity was measured using the Zen software. All imaging and analyses were conducted by a blinded investigator.

### Four-Limb Hang Test

Whole body endurance was measured using a modified version of a previously established protocol (76). Mice were suspended upside down on a rigid wire grid at a height of 30 cm. Hang time prior to falling onto a padded surface was measured across five trials for each mouse, with a five-minute break between trials. The hang impulse was calculated as the average time of the three median trials multiplied by mouse body weight. For injured animals, hang tests were performed the day before injury and repeated again immediately prior to injury, and at 1, 7 and 14 days post injury. For uninjured animals, hang tests were conducted prior to AAV injection, then at 1, 7, and 14 days after injection.

### In Situ Contractile Testing

TA strength was evaluated using an *in situ* testing apparatus (Model 809B, Aurora Scientific, Aurora, ON, Canada). The hindlimb skin of anesthetized mice was cut to expose the peroneal nerve, and the Achilles tendon was severed to prevent a contribution of the muscle antagonist. The foot was secured onto a footplate positioned at 20° of plantar flexion. Initially, single twitch stimulations were elicited to measure the peak twitch, time to peak twitch and the half-relaxation time. The muscles then received 350-ms train tetanic stimulations at different frequencies with two-minute intervals between contractions. Animals were then given a recovery period of 10 minutes before proceeding to the high frequency fatiguing protocol wherein the muscles were activated by a stimulus train at 100 Hz every four seconds for seven minutes. Recovery was measured five and ten minutes after cessation of the fatigue protocol. Force values were normalized to muscle cross-sectional area (CSA) as previously described(77). Animals were euthanized immediately following testing via cardiac puncture, and blood and tissues were collected for analyses.

### Circulating Klotho and FGF23 quantification

Blood collected from experimental and control animals was clotted at room temperature for one hour, after which it was spun at 16,100 G for 15 minutes in a fixed rotor centrifuge at 4°C in order to separate serum. This serum was diluted 1:25 and 1:6 in diluent buffer to quantify Klotho (Cloud Clone Corp), and FGF23 (abcam), respectively using commercially available ELISA kits.

### AAV production

Recombinant AAV8 was used as vector for expressing murine Klotho or enhanced green fluorescent protein (GFP). The genomic constructs contained the LP-1 promotor, GFP or a codon-usage optimized sequence of murine alpha klotho (Geneart, Regensburg, Germany) with an N-terminal secretion signal originating from human CD33 (sequence: MPLLLLLPLLWAGALA) and a C-terminal V5-tag epitope (78), followed by a WPRE sequence and an SV40 polyA signal. The expression cassettes were flanked by AAV2-derived inverted terminal repeats. The recombinant AAV8-LP1-mKlotho and AAV8-LP1-eGFP vectors were prepared as previously described (79). Briefly, human embryonic kidney cells (HEK-293H cells, Thermo Fisher Scientific) were cultured in Dulbecco’s modified Eagle’s medium + GlutaMAX-I + 10% fetal calf serum (Gibco/Thermo Fisher Scientific) and transfected as previously described (80). AAV purification via polyethylenglycol precipitation, iodixanol gradient, ultrafiltration and sterile filtering was conducted as described previously (79). Genomic titers of purified AAV8 vector stocks were determined by isolation of viral DNA and subsequent qPCR analysis using primers specific for the LP-1 promotor with the following primer sequences; forward: GACCCCCTAAAATGGGCAAA. reverse: TGCCCCAGCTCCAAGGT.

### AAV-Administration, Injury, and Sarcopenia Experimental Models

Animals were restrained using a tail illuminator restrainer (Braintree Scientific Inc.) and received a tail vein injection of either AAV-Kl or AAV-GFP. Solutions were diluted in Dulbecco’s phosphate-buffered saline to a final volume of 100 µL/mouse. For the muscle regeneration experiment, the dose of GFP and AAV-Kl used was 1×10^10^ vector genomes(vg) per animal. Five days following injection, each mouse received a bilateral intramuscular injury to the tibialis anterior muscle (TA) using 10 µL of 1 mg/mL cardiotoxin (*Naja pallida*, Sigma, molecular weight 6827.4), and animals were euthanized 14 days after injury. In uninjured animals, the dose used was 3×10^8^ vector genomes per mouse, and mice were euthanized 14 days after AAV injection.

### RNA seq Analysis

We performed RNA sequencing on gastrocnemius muscle collected from female mice injected with AAV-GFP and AAV-Kl. Four animals were used for each of the five groups: young GFP, old GFP, old Klotho, oldest-old GFP, and oldest-old Klotho. Raw fastq files were converted to gene counts after filtering and alignment using Partek Flow software. The visualizations were generated using R packages, ggplot2, clusterprofiler, VennDiagram and igraph. DE analysis was performed with Deseq2. DE genes with P value less than 0.05 were defined as significantly different.

To identify genes associated with the hallmarks of aging, a list of 77,708 genes derived from Mouse Genome Informatics’ batch query was generated based on the GO terms related to cell biology of aging **(Supplemental methods)**. The ratio of DE genes that changed under each hallmark was visualized in the barplot and then network plot. Shannon network entropy was computed after mapping genes to proteins using Biomart in R **(Supplemental methods)**. Protein-protein interaction (PPI) network was obtained from STRING database v11.0 (45).

## Acknowledgement

The authors thank Matthias Duechs from Boehringer-Ingelheim (BI), who performed MSD-ELISA on samples obtained from the dose testing experiment in young animals. We thank Timothy Lezon, Assistant Professor, Computational and Systems Biology at the University of Pittsburgh for helping us understand and interpret the concept of entropy. We also thank Center for Biologic Imaging, University of Pittsburgh for providing resources to perform histological IF staining. The studies reported in this manuscript are supported by funding from NIA R01AG052978 (F.A.), NIA R01AG061005 (F.A.), and BI (F.A.).

