## Supplemental figures, methods, tables for "The biphasic and age-dependent impact of Klotho on hallmarks of aging and skeletal muscle function"

Supplementary Figure 1

**|____*____|**

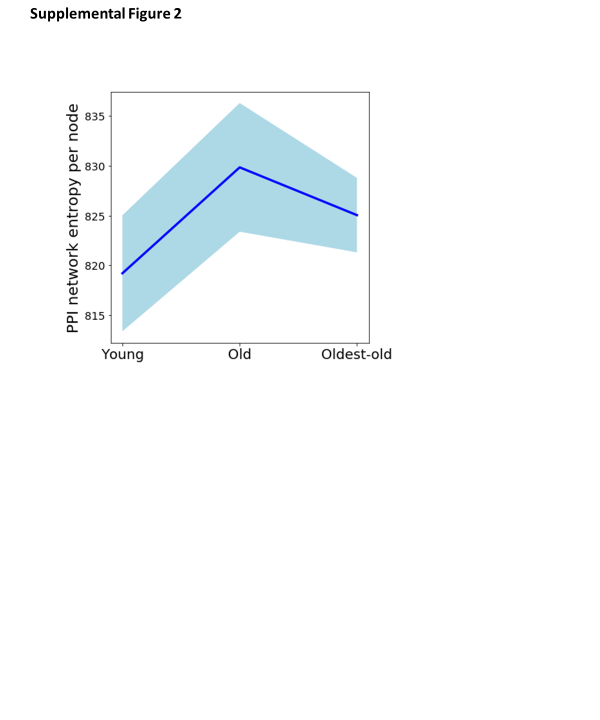

**S1. Network entropy trend with all genes considered in the PPI network.** Protein-protein interaction (PPI) network entropy computed from transcriptomic data indicates an increase in transcriptional noise of genes from young to old, after which time entropy declines slightly. A non-parametric Kruskal Wallis test and Dunn’s post-hoc test was performed (*p = 0.056).

Supplemental Figure 2

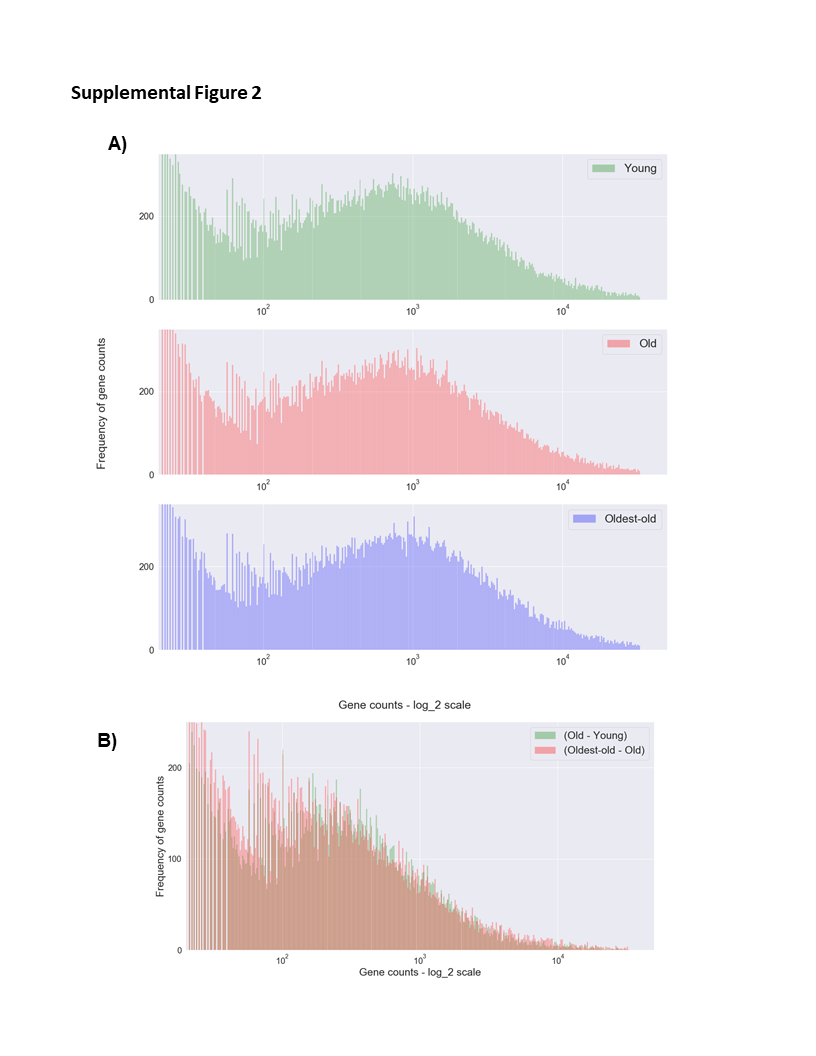

**S2. Histogram of all gene counts across young, old and oldest-old age groups.** (A) Gene counts concatenated values from 4 animals from each age group, young, old and oldest-old. (B) Histogram that shows the difference between age groups, with the green (Old - Young) histogram right shifted compared to red (Oldest-old – Old).

Supplemental Figure 3

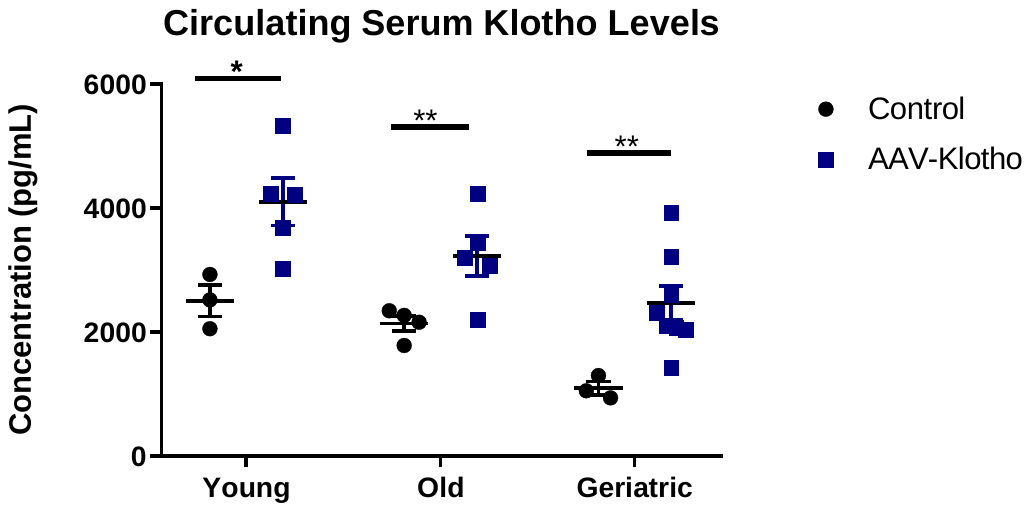

**S3.** **AAV-Klotho increases circulating levels of Klotho in uninjured female mice**. Circulating Klotho levels two weeks post AAV-Klotho injection (3x10^8^ vg/ml) compared to untreated controls (One-tailed Mann-Whitney U test, Experimental AAV-Klotho groups compared to age- and sex-matched controls, *p<0.05, **p<0.01, n=3-8/group). Data presented as mean ± SEM.

Supplemental Figure 4

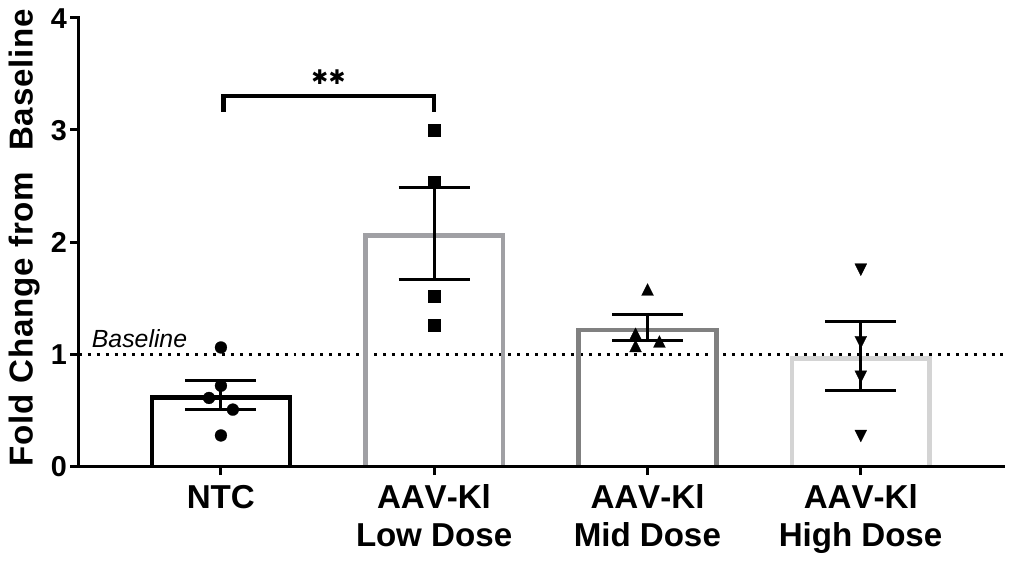

**S4.** **Dose testing of AAV-Klotho administration on whole-body endurance.** The ratio of whole-body endurance of old (21-24 months) female mice 14-days after receiving AAV-GFP or AAV-Kl at three different doses (3x10^8^ vg/mouse, 1x10^9^ vg/mouse, and 3x10^9^ vg/mouse) (one-way ANOVA, **p<.01, n=4-5/group). Data presented as mean ± SEM.

Supplemental Figure 5

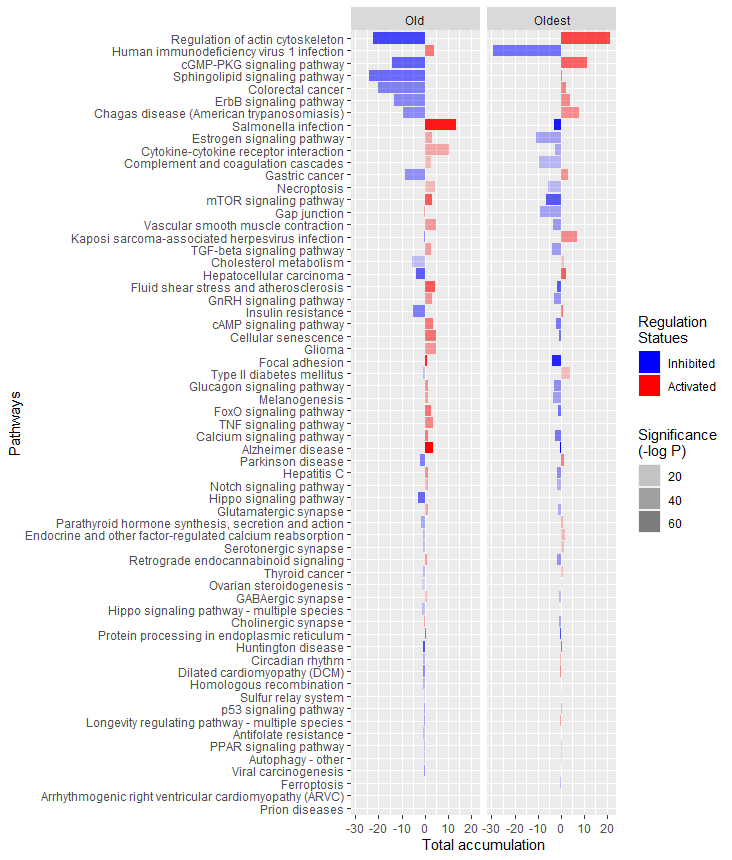

**S5.** **Signaling Pathway Impact Analysis (SPIA) showing all affected KEGG pathways.** Bar plot showing all the KEGG pathways that change in opposing fashion between old and oldest-old groups after AAV-Kl treatment ranked by largest absolute difference in total accumulation.

Supplemental Table 1

**Table S1: Antibody List.** Primary antibodies and corresponding dilutions used for immunofluorescence imaging.

| **Antibody** | **Source** | **Dilution (in .1% Triton-X + 3% BSA+ 5% goat serum in PBS)** |
| --- | --- | --- |
| Rabbit anti-Laminin | Abcam ab11575 | 1:500 |
| Lipidtox Red | Invitrogen H34476 | 1:200 |
| Collagen IV | Abcam ab6586 | 1:500 |

Supplemental Table S2

**Table S2. Contrasting gene response to AAV-Kl administration between age groups.** Table displaying the biphasic gene expression responses in the top three KEGG pathways (**Figure 6D**) between old and oldest-old female mice after AAV-Kl treatment.

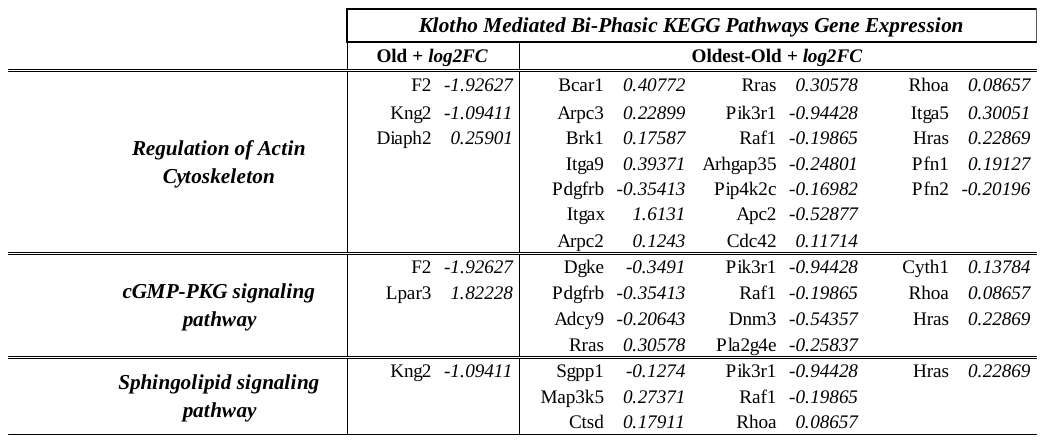

**SUPPLEMENTAL METHODS**

***Study rigor***

All animal experiments were approved by the University of Pittsburgh’s Institutional Animal Care and Use Committee. Experiments included young (3-6 months), middle-aged (10-14 months), old (21-24 months), and oldest-old (27-29 months) male and female C57 BL/6J mice, obtained from Jackson Laboratories, with the exception of AAV dose testing animals which were obtained from Charles River Laboratories. All mice were housed in pathogen free conditions under a 12 hour12-hour light and 12- hour dark cycle. Virus exposed animals injected with AAV were housed and experimented on under higher biosafety regulations in the animal facility (Biosafety level II). Animals with obvious health problems (i.e, tumors, malocclusion, etc.) were eliminated prior to inclusion into the study where possible and excluded from endpoint analyses if pathologies were noted during experimentation. Given the large amount of variability in physical endurance capacity, animals were randomized into cohorts only if they met the criteria of falling into the 25th-75th percentile for the four-limb hang test (described below). All animals meeting criteria for inclusion were randomized to treatment groups and acclimatized for at least one week. For *in vivo* experiments, each experiment was repeated across a minimum of two cohorts, and experimenters performing endpoint analysis were blinded to the experimental groups. Where possible, power analysis from pilot data were done to select the number of animals needed for the study. Power analyses were performed *a priori* assuming a two-sided alternative hypothesis with alpha=0.05 and 80% power (G*Power 3.1.9.2). Based on the prior findings from the laboratory on peak tetanic force in young and aged animals and after adjusting for unforeseen circumstances estimated at 20%, the final sample size was calculated to be 8-10 animals/group.

***Cryopreservation and preparation of skeletal muscle tissue***

Post euthanasia (see serum collection below), the tibialis anterior (TA) muscles of mice were carefully extracted and wet weight was measured using a standard balance. Immediately after excision, TAs were frozen in liquid nitrogen cooled 2-methyl butane for one minute, and stored at -80°C. Slides for histological analyses were prepared using the Thermo Fisher CryoStarNX50 cryostat. Tissue was cut at 10μm with the cryostat set at -25°C. Each slide (up to 20) captured equidistant tissue sections throughout the entire length of the TA. Slides were stored in the -80°C freezer until use for staining.

***Immunofluorescence Imaging***

One slide from each experimental replicate was thawed and immediately fixed by covering sections with 2% paraformaldehyde using a pipette for 10 minutes. After fixing, slides were washed with 1X-Phosphate Buffered Saline (PBS) three times for two minutes each. A hydrophobic barrier was then drawn around the sections, and they were permeabilized with 0.1% triton-X (Fluka 93420) in PBS for 15 minutes, followed by a one-hour blocking step using 0.1% triton-X plus 3% Bovine Serum Albumin (BSA, Sigma A7906) in PBS. Primary antibodies were diluted (**Table S1**) in a solution of 0.1% triton-X, 3% BSA, and 5% normal goat serum in PBS then added onto the sections and incubated overnight at 4˚C. One negative control slide per staining set was generated by deleting the primary antibody in the antibody solution. The secondary antibodies used were goat anti-rabbit and goat anti-rat IgG cross-adsorbed antibodies (Invitrogen) and were diluted in the same base solution as the primary antibodies. After washing three times for two minutes in PBS, the secondary antibodies were added to the sections for one hour and incubated in the dark at room temperature for one hour. The slides were washed with PBS three times and DAPI (1:500 dilution in PBS) was added onto the slides for two minutes. Next, slides were washed twice with PBS, dried and mounted with coverslips using Gelvatol mounting medium (Source: Center of Biologic Imaging, University of Pittsburgh). Slides were allowed to dry for at least 24 hours at 4˚C prior to imaging.

Slides were imaged using a Zeiss Observer Z1 semi-confocal microscope. For uninjured slides, regions of interest were randomly selected by navigating to an imaging site of normal nuclear density based on the DAPI channel and capturing an image. At least three images were captured per slide (each slide contained tissues from one animal). All images were collected at 20x magnification. Negative control slides were used to threshold for the signal intensity and to set the exposure time for individual channels. The exposure time was kept consistent for each set of stains based on the negative control. For injured tissues, the injury site was located by finding the localized area of high nuclear infiltration-a known sign of maximal injury. Images were then collected at the injury site. At least three images were collected per slide. Intensity was analyzed for Collagen IV and lipid using the Zen software and was normalized to image area. Myofiber CSA was quantified from 20X images using Image J software. Rabbit anti-Laminin was used to define myofiber borders and an auto-threshold mask was used to calculate the average myofiber area across all collected images per mouse. Fibers displaying faded laminin staining and/or fibers extending into the edges of the image, were excluded from analysis (1).

***Four-limb hang test***

The four-limb hang test was used to measure mouse muscular endurance non-invasively and, thus, characterize performance before and/or after pharmacological administration. Mice were suspended upside down from a steel mesh grid (1cm x 1cm squares) above a custom-built chamber (~30 cm high) with appropriate padding to prevent harm. Four limb strength was evaluated using a Hang Impulse (HI) score (*bodyweight in grams* **x** *time spent hanging*). The time spent hanging was recorded from the time of inversion until all paws are released from the grid. Prior to recorded trials, mice were acclimatized to the grid and the chamber for five minutes. Mice are subject to five trials in one session with a gap of at least 5 minutes between trials. Sessions were conducted at roughly the same time of the day (between 10-12 pm) and in the same progression across trials (i.e. the first mouse on session one would also be first on session two). Evaluation of performance is calculated as the average HI of three best trials. Baseline endurance capacity was evaluated prior to experimentation.

For muscle regeneration experiments, injured animals treated with AAV-GFP or AAV-Kl, hang tests were performed before and after injections, and repeated immediately prior to injury, and then again at one, seven and 14 days post injury. For uninjured animals, hang tests were conducted prior to AAV injection, then at one, seven, and 14 days after injection.

***In situ contractile testing***

TA muscle strength of animals in the aged cohorts and treatment cohorts was evaluated using an *in situ* testing apparatus (Model 809B, Aurora Scientific, Aurora, ON, Canada). Anesthetized mice underwent surgery to expose the peroneal nerve for stimulation and the Achilles tendon was severed to prevent a counteracting response. The foot was secured onto a footplate in 20° of plantar flexion for maximal responses(2). Initially, single twitch stimulation tests were conducted to measure the peak twitch, time to peak twitch and the half-relaxation time. Shortly after, tetanic stimulations (350-ms train) at 10, 30, 50, 80, 100, 120, 150, 180 and 200 HZ with two-minute intervals between contractions were induced to obtain the force-frequency curves. Post tetanic stimulation, animals were given a recovery period of 10 minutes before proceeding to the high frequency fatiguing protocol wherein muscles were activated every four seconds by a stimulus train at 100 Hz for 7 minutes. After completion, force recovery was evaluated at 5 and 10-minute timepoints. Results of both the single and tetanic stimulations were collected in torque (mN-m) and the absolute force values (mN) were extrapolated by dividing by the length of the footplate. Values were then normalized to muscle cross-sectional area (CSA) as described by Brooks and Faulkner (3). Results of the high frequency fatigue and recovery test were analyzed as percentage values of the initial response.

***Hallmarks of aging genes classification***

First, we compiled a list of key functions and changes for all systemic hallmarks of aging and cellular hallmarks of aging (4-7). Next, we created multiple gene lists by searching for different combination of gene ontology (GO) terms on the MGI GO search database (See bullet points below). A table of (genes) x (hallmarks) was formed by genes classified into different hallmarks of aging. Note that, GO terms associated with multiple hallmarks of aging were accounted for in this process.

From this step onwards, we focused on each hallmark of aging, as opposed to a specific gene or a GO term. The goal of the study was to identify and target the genes associated with the hallmarks of aging, to reverse sarcopenia.

Important points about [MGI GO search](http://www.informatics.jax.org/marker): (Used batch query forms)

- AND – together appear
- OR – default with space (only one or both)
- NOT-
- ANDNOT
- If 2 words together make 1 word, you use quotations.
- Comma is ignored, use quotation if the term itself contains ‘,’

Hallmarks of Aging

1. Genomic Instability
2. Telomere attrition
3. Epigenetic alteration
4. Loss of proteostasis
5. Nutrient sensing deregulation
6. Mitochondrial dysfunction
7. Cellular senescence
8. Stem cell exhaustion
9. Altered intracellular communication

Cellular Hallmarks of Aging

1. Replicative aging, senescence and renewal (#7)
2. Chromosome and telomere attrition (#1, #2)
3. Transcriptional regulation (#3)
4. Nuclear trafficking and organization (#1)
5. Protein translation (#4)
6. Proteostasis (#4)
7. Unfolded protein response (#4)
8. Autophagy (#4)
9. Mitochondrial function, biogenesis, mitophagy (#6)
10. Cytoskeletal integrity (#6, #9)
11. Cell membrane and ECM (#9)

Breaking down hallmarks:

1. Genomic Instability

- Somatic mutations of nuclear DNA
- Mutations and deletions in aged mtDNA
- Defects in nuclear lamina
- DNA repair mechanisms
  - “DNA stability” (4 genes)
  - “nuclear lamina” (13 genes)
  - “Mitochondrial DNA” (26 genes)
  - “DNA repair” (495 genes)

2. Telomere attrition

- “telomere” (193 genes)
- ”telomerase” (96genes)
- “ROS” (430genes)

3. Epigenetic alteration

- Histone modifications
- Chromatin remodeling
- DNA methylation
- Transcriptional alterations
- Reverse of epigenetic changes
  - “histone modification” (517 genes)
  - “post-translational_modification” (25 genes)
  - “chromatin remodeling” (181 genes)
  - “DNA methylation” (106 genes)

4. Loss of proteostasis

- Chaperone-mediated protein folding and stability
- Proteolytic systems
- “proteolytic” (66 genes)
- “autophagy” (403 genes)
- “proteasome” (943 genes)
- “heat-shock protein” (251 genes)
- “ubiquitin” (1324 genes)
- “protein translation” (617 genes)
- “unfolded protein response” (67 genes)

5. Nutrient sensing (NS) deregulation

- Insulin and IGF-1 signalling pathway
- Other NS systems – mTOR, AMPK, sirtuins
- “insulin” (572 genes)
- “growth factor” (972 genes)
- “mTOR” (14 genes)
- “AMPK” (34genes)
- “identical protein binding” AND “transcription factor binding” (168 genes) *-> sirtuin is associated with these GO terms*
- “NAD” (469 genes)
- “glucose metabolism” (203 genes)
- “oxidative metabolism” (286 genes)

6. Mitochondrial dysfunction

- Reactive oxygen species (ROS)
- Mitochondrial integrity and biogenesis
- Mitohormesis
- “mitochondria” (1961 genes)
- “ROS” (430 genes)
- “mitophagy” (67 genes)

7. Cellular senescence

- Non-telomeric DNA damage
- Derepression of the INK4/ARF locus
- Senescence-Associated Beta Galactosidase(SABG)
- Excessive mitogenic signaling
- p16Ink4a/rb and p19ARF/p53 pathways – relate to senescence and oncogenic insults (cell cycle inhibitory proteins)
- Senescence-Associated Secretory Phenotype (SASP) – explains how senescent cells alter tissue microenvironment
  - “senescence” OR “senescent” (74 genes)
  - “mitogen” (841 genes)

8. Altered intracellular communication

- Inflammaging
- Increased activation of NFkB TF
- Altered autophagy response
- Increased levels of IL-1b, TNF, and interferons
- Decay factor AU-binding factor 1
- *include ECM changes*
- Cytoskeletal changes and aging
- Cell membrane, ECM and aging
  - “extracellular matrix” (829 genes)
  - “extracellular vesicle” (105 genes)
  - “paracrine” (221 genes)
  - “interferon” (1198 genes)

9. Stem cell exhaustion

- Diminished production of adaptive immune cells (immunosenescence)
- Gtpase cdc42 activity is increased in aged hematopoetic stem cells

This hallmark was not included in our final analysis because all the keywords for stem cell exhaustion were already covered under in one of the other hallmarks. This would have resulted in no unique genes for “stem cell exhaustion”.

***Network entropy methodology and interpretation***

Pre-processing RNA-seq to protein-protein interaction (PPI) networks

Raw gene count matrix was used to generate PPI networks. For each sample, we generated a node-list and an edge-list. Absolute gene expression values were used to create the node-list. Edge-list was generated with mouse PPI networks obtained from STRING database version 11.0. If the gene counts were zero, then the nodes were not considered as a part of the PPI network. This process was done for all the genes, as well as only for the genes classified into hallmarks of aging.

Network entropy interpretation

In the field of information theory, Shannon entropy is the measure of surprise of an outcome. For example, a fair coin two has two equally probable outcomes. This would be the case of maximum surprise, or maximum Shannon entropy. On the other hand, if the coin is rigged, and probability of obtaining heads is 0.9, then one will not be surprised to get heads. This is the case of lower Shannon entropy. When this concept is extrapolated to graph theory, we aimed to predict the presence or absence of an edge in the graph. If network entropy is high, this means that it is harder to predict the network connections with the knowledge we have. A higher network entropy is bound to happen when the knowledge you have about the system is insufficient or highly variable.

The principle of maximum entropy states that the probability distribution best representing the current state of knowledge on a system is the one which maximizes the Shannon entropy, given the prior knowledge about the system (8). Using this method, we gain maximally unbiased information in the absence of complete knowledge. In the content of network biology, the ensemble of most probable network configurations are the ones that have maximum entropy, given certain constraints.

Our goal was to capture the change in gene expression distribution across age groups using network entropy. In this study, we obtained nodelists from the RNA-seq experiment, and edgelists obtained from existing databases (eg. STRING) that have PPI network. In addition, we accounted for relative gene expressions by weighting the edges of the network to incorporate the current knowledge we have about the PPI network connections. Hence, we hypothesized that changes in gene expressions will affect the number of nodes, number of edges and the edge weights–a phenomenon that could potentially be captured by the network entropy metric.

Conventionally, maximum entropy is used to predict the probability distribution. But in this study, we used network entropy values corresponding to the most probable ensemble of PPI networks. This allowed us to quantify the change in transcriptional noise and variation in gene expressions, into a single integrative metric.

Network entropy calculation

Network entropy is a constraint-based optimization problem, that is solved using Lagrangian multipliers. Objective function to maximize is the Shannon entropy of the network. A value of network entropy will correspond to a probability distribution for edges. There are two constraints for the optimization problem. The first constraint imposes a degree sequence of the network, whereas the second constraint imposes restriction on the edge weights of the network. The concept is explained with an arbitrary simple example, as shown below.

| Gene counts for node-list (15 genes) | | | | | | | | | | | | | | | Age and # nodes (n) |
| --- | --- | --- | --- | --- | --- | --- | --- | --- | --- | --- | --- | --- | --- | --- | --- |
| 500 | 700 | 0 | 0 | 0 | 0 | 0 | 100 | 58 | 350 | 0 | 675 | 10 | 509 | 0 | Young (n=8) |
| 502 | 704 | 0 | 1 | 2 | 3 | 0 | 95 | 59 | 375 | 0 | 450 | 0 | 458 | 0 | Young (n=10) |
| 600 | 690 | 0 | 1 | 0 | 5 | 0 | 110 | 65 | 378 | 2 | 567 | 9 | 450 | 0 | Old (n=11) |
| 595 | 721 | 0 | 0 | 0 | 9 | 0 | 100 | 0 | 300 | 0 | 604 | 0 | 425 | 0 | Old (n=7) |

Let edge-list is fixed: (1-3), (3-7), (5-8), (8-9), (9-12), (10-14)

Constraint 1:

The degree sequence was fixed based on the edge-lists. Zeroes are not considered in the network as nodes.

Constraint 2:

The edge weights were calculated by taking the difference between gene counts between the connected nodes. To normalize for different number of nodes in the network, the edge weights are binned with number of bins equal to square root of the number of nodes in the network.

There would be sqrt(n) ~3 bins. [0-240, 241- 482, 483-721]

The binning distribution will vary according to the relative gene expression changes. If the binning is similar for all the samples, then network entropy values will be very close to each other. If the binning is different, network entropy values will be different.

***Serum Collection***

Animals were anesthetized using isoflurane and remained anesthetized throughout the duration of serum collection. The animal was placed in a supine position and its paws were taped down to spread out the torso. A small incision was made with scissors and forceps in the skin above the xiphoid process of the sternum and expanded to reveal the bottom of the rib cage. The peritoneum in the same area was then cut to reveal the underside of the diaphragm. The diaphragm was cut and cleared away with scissors, thus exposing the bottom of the heart. A clamp was used to hold the ribcage out of the way, and a 25 ½ G needle attached to a 1 mL insulin syringe was used to collect blood directly from the apex of the heart. Using this method, roughly 800 µL of blood could be collected from each animal. After collection, the animals were euthanized via cervical dislocation. The collected blood was allowed to clot in a 2 mL tube at room temperature for one-hour. Then, the blood was spun at 16,100g for 15 minutes at 4˚C in a microcentrifuge. The serum was collected and aliquoted into 50 µL tubes and stored at -20˚C until used.

***ELISA***

Klotho ELISA

The Klotho ELISA was conducted according to the protocol using the Cloud Clone Corp. ELISA Kit for mouse Klotho (SHE757Mu, Lot: L180223640). This kit was validated by Sahu et al (9). Briefly, blood serum samples and standards were diluted in PBS (1:25) and 100 µL of the samples were added in duplicates to a 96-well plate pre-coated with a biotin-conjugated antibody specific for Klotho detection. This plate was then incubated at 37°C for one hour. Samples and standards were removed and 100 µL of biotin-conjugated antibody cocktail was added to each well and incubated at 37°C for one hour. The microplates were washed three times with washing buffer provided by the kit (two minutes per wash), using an automated plate washer (BioTek 50TS). Following this, 100 µL of Avidin conjugated Horseradish-Peroxidase was added to each well and incubated at 37°C for 30 minutes. The plate was washed with buffer five times, following which 90 µL of tetramethylbenzidine (TMB) substrate was added to each well and the plate was incubated at 37°C for 20 minutes. A sulfuric acid-based stop solution was added to terminate the reaction and optical density of each well was measured with a Spectramax M3 plate reader (Molecular Devices) at wavelength of 450 nm. The analysis was conducted using Microsoft Excel by generating a 2P equation for concentration based on the standard curve and plotting the averaged absorbance values for each sample along this curve.

Any sample displaying evidence of hemolysis were not included in data analysis, as we have seen that hemolyzed interferes with values. Samples were not subjected to multiple free-thaw cycles, as we have observed that this may affect α-Klotho detection (10).

FGF23 ELISA

The FGF23 ELISA was performed according the protocol provided by the manufacturer (Abcam, ab213863, Lot: GR3326863). Briefly, blood serum samples and standards (diluted 1:6 in sample diluent buffer) were added in duplicates to a 96-well plate pre-coated with antibody against FGF23 and incubated for 90 minutes at 37°C. The plate content was then discarded and the plate was blotted. Following this, a biotinylated anti-mouse FGF32 antibody was added to all wells and incubated for 60 minutes at 37°C. Next, the plate was washed three times with 0.01M PBS using the same plate washer as described above. 100 µL of avitin-biotin-peroxidase complex was added to each well and incubated again for 30 minutes at 37°C. The plate was washed five more times, following which 90 µL of TMB substrate was added to each well and incubated in the dark for 20 minutes at 37°C. Finally, 100 µL of TMB stop solution was added and the plate was immediately read at 450 nm using the same plate reader as described above. Analysis was conducted as described above. Hemolyzed samples were not included, and each sample was frozen once prior to the experiment.

Meso-Scale Discovery (MSD) Klotho ELISA

Detection of circulating Klotho levels from mouse blood samples was performed by using an ELISA assay on the MSD platform. Standard-bind MSD plates (#L15XA-1, MSD) were coated with mouse Klotho capture antibody (#AF1819, R&D) at a final concentration of 4 µg/mL in PBS for 1 hour at room temperature. Subsequently plates were washed three times using wash buffer (PBS + 0.05% Tween-20 ) at 300 µL/well followed by an incubation with 3% blocker A solution (R93BA-2, MSD) for 1 hour at room temperature. Dilutions of serum samples and murine recombinant Klotho standard (#1819-KL-050, R&D) were prepared in 1% blocker A solution and added to the MSD plate in a final volume of 25 µL/well after the plate was washed for three times. Samples were incubated for 1 hour at room temperature followed by three washing steps. Detection antibody (#BAF1819, R&D) was diluted to 1 µg/mL in 1 % Blocker A solution in PBS and SULFO-tag labelled streptavidin (#R32AD-5, MSD) was diluted to 0.5 µg/mL in 1% Blocker A in PBS. Both dilutions were added to the plate simultaneously (25 µL/well each) and incubated for 1 hour at room temperature. After three washing steps, 1x Read Buffer (#R92TC-2, MSD) diluted in water was added at 150 µL/well. Electrochemiluminescence was detected in the MSD Sector Imager 600.

***Metabolite Analysis***

Serum metabolites (ie. glucose, cholesterol and free lipids) from blood serum were analyzed through a third-party vendor, *Charles Rivers Biomarker Services: Clinical Pathology and Immunology*.

***AAV Tail Vein Injection***

Aliquots of AAV-Kl (1.79e13 Vg/ml) and AAV-GFP (3.48e12 Vg/ml) were stored in -80°C and thawed within an hour of use. Animals were restrained using a tail illuminator restrainer from Braintree Scientific Inc. and injected with either a GFP expressing non-targeting control or AAV-Kl via the tail vein using a 31G insulin syringe. Viral solutions were diluted in Dulbecco’s phosphate-buffered saline (DPBS) to a final volume of 100 µL/mouse. A successful injection entailed full administration of the viral solution into the vein smoothly, without swelling and bleeding following syringe removal. Only successfully injected mice were included in the study (a total of six were excluded). For the muscle regeneration experiment, the dose of GFP and AAV-Kl used was 1x10^10^ vector genomes (vg) per animal. Five days following injection, each mouse received a bilateral intramuscular injury to the tibialis anterior muscle (TA) using 10 µL of 1 mg/mL cardiotoxin (*Naja pallida,* Sigma, molecular weight 6827.4). Animals were euthanized 14 days after injury. For uninjured animals, the dose used was 3x10^8^ vg per mouse and animals were euthanized 14 days post injection.

***AAV-Klotho in vivo dose response study***

Female C57Bl/6 mice (9–12 week old) with a body weight of 19–21 g were purchased from Charles River Laboratories. AAVs were diluted to the desired concentrations in PBS and administered into the tail vein under light isoflurane anesthesia. The final volume for injection was 100 μL per mouse. Three weeks after injection with AAV-GFP, AAV-Kl or DPBS, animals were sacrificed and blood samples were collected for the detection of circulating Klotho levels by MSD-ELISA. Animal experiments in this study were approved by the local German authorities (Regierungs-präsidium Tübingen) and conducted in compliance with the German and European Animal Welfare Acts.

***Statistical Analysis***

Analyses were performed using GraphPad Prism version 8 software. Shapiro-Wilk and Levene’s tests were initially performed to assess normality of data and equality of variances, respectively. If assumptions of normality and homogeneity of variances were met, a Student’s t-test was performed while comparing two groups. When conditions for normality were not met, the groups were compared using Mann-Whitney U test. A Welch’s test was applied when there were differences between the standard deviations of the groups. All results were expressed as mean ± standard error. Statistical significance was established, a priori**,** at *p*≤0.05.

**Supplemental** **References**

1.	Mula J, Lee JD, Liu F, Yang L, Peterson CA. Automated image analysis of skeletal muscle fiber cross-sectional area. J Appl Physiol (1985). 2013;114(1):148-55.

2.	Distefano G, Ferrari RJ, Weiss C, Deasy BM, Boninger ML, Fitzgerald GK, et al. Neuromuscular electrical stimulation as a method to maximize the beneficial effects of muscle stem cells transplanted into dystrophic skeletal muscle. PLoS One.8(3):e54922.
